## Supplementary figures and images for "Boldine alters serum lipidomic signatures after acute spinal cord transection in male mice"

### Fig. S1

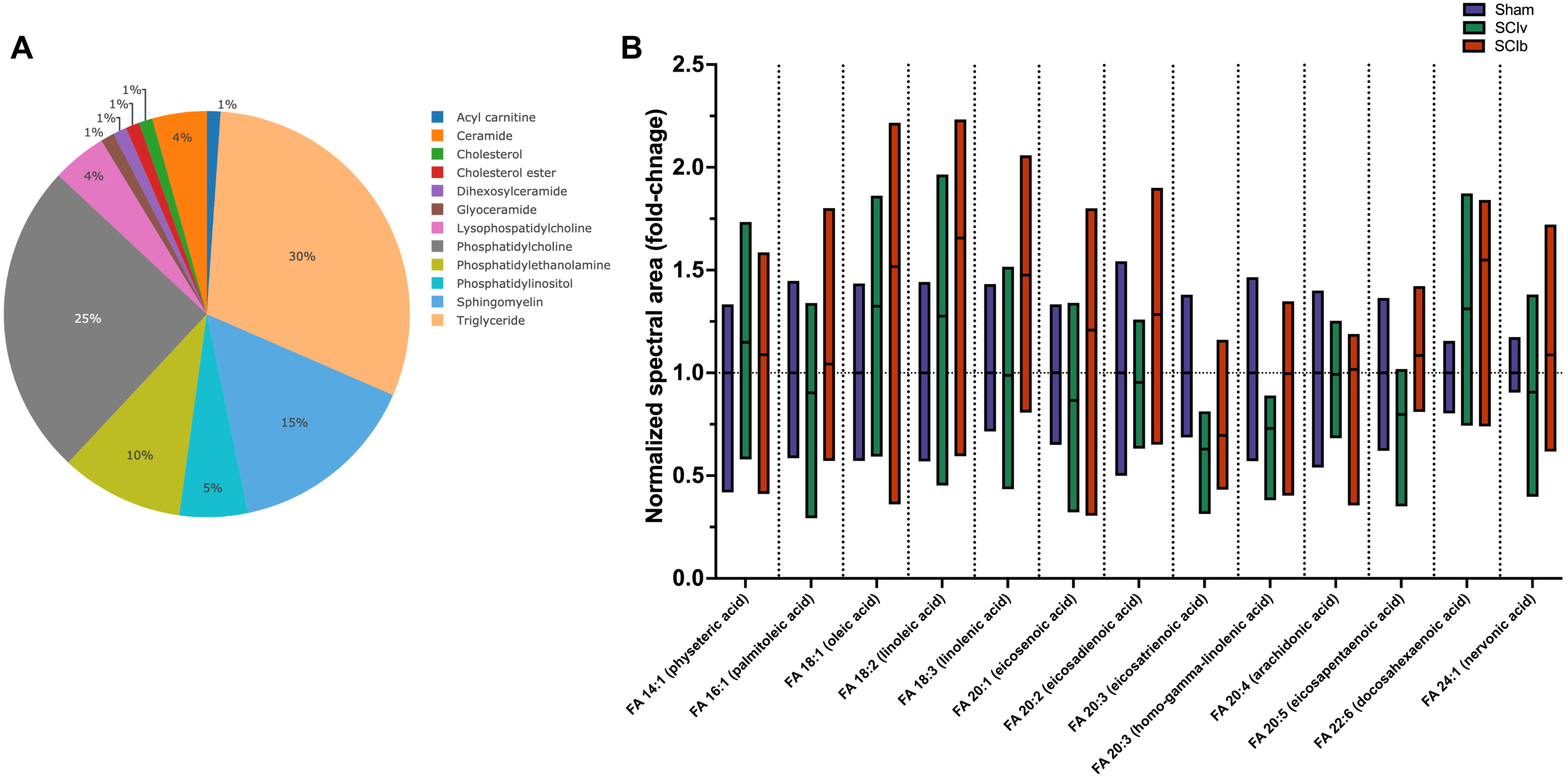
